# Environmental programming of adult foraging behavior in *C. elegans*

**DOI:** 10.1101/400754

**Authors:** Sreeparna Pradhan, Sabrina Quilez, Kai Homer, Michael Hendricks

## Abstract

Foraging strategies must be tuned to the availability and distribution of resources in the environment. This can occur over generations and lead to genetic differences in foraging behavior, or it can occur on shorter time scales within an individual’s life span. Both genetic and experience-based strategies must be implemented by neural circuits that respond to environmental cues and track internal states, and the analysis of such circuits provides insight into the neural basis of complex decision making. In C. elegans, between-strain genetic differences and within-strain plasticity in foraging has been observed. Most individual changes in foraging are short-term, based on experience over several hours. Here, we tested if developmental experience could permanently alter foraging. We found that in wild strains that are normally highly exploratory, early-life starvation leads to “cautious” foraging behavior in which exploration is reduced. We characterize the behavioral bases for these strategies and identify changes in the dynamics of a locomotory circuit involved in navigation. Overall, we show that some C. elegans strains exhibit adaptive tuning of their foraging behavior based on early-life experience, and this is associated with changes in a core navigation circuit.

## Introduction

Animal behavior is an outcome of a complex interplay of inherent genetic traits, immediate sensory experience, memories of past experience, and current internal states like nutrition and motivation. Experience-dependent plasticity in behavior can act over different time scales, and early life experiences can be especially crucial in regulating long-term changes in adult behavior. A large body of work in ethology described how experiences during a critical period in development can permanently alter adult behavior in many species of birds, fishes and insects (Bateson, 1966; Hess, 1959). The most famous of these studies is perhaps Konrad Lorenz’s work on geese, where goslings “imprint” on the first moving object they see and form an attachment towards it (Lorenz, 1937). Many imprinted behaviors such as food preferences, selection of a territory, and mating choices have a strong adaptive role in an ecological context (Immelmann, 1975).

Adverse environmental conditions in early life can also affect adult health, in both animals and humans. Rodents and nonhuman primates receiving low maternal care have been shown to develop heightened stress responses in adulthood (Coplan et al., 1996; Hales and Barker, 2001; Ladd et al., 2000). Growing evidence from human epidemiological studies have linked poor maternal diet, childhood malnutrition and early life stress to adult onset metabolic and psychiatric disorders (Hales and Barker, 2001; Painter et al., 2008; Song et al., 2009). Many such disorders have complex origins, where early life environmental stress couples with genetic predispositions to affect behavioral plasticity in adults. A model for adaptive tuning of behavior suggests that behavioral modifications depend on environmental fluctuations and trait plasticity (Sih, 2011). In response to changing environmental conditions, behavioral plasticity is adaptive as animals tune behavior to a predicted environment. A stable environment favors low trait plasticity and fluctuating environments favor higher trait plasticity. While a number of studies have identified behavioral changes in response to early life environmental stress, it is less clear how these stressful conditions act in the context of different genetic backgrounds to produce different behaviors.

The nematode *Caenorhabditis elegans* goes through four larval stages (L1 to L4) to reach adulthood in the presence of plentiful food (well-fed). Environmental stress (starvation, high population density, high temperature) in the late L1 stage triggers entry into a developmental arrest phase called dauer (Golden and Riddle, 1984, 1982). Dauer larvae can endure stressful conditions for many months. When conditions improve, larvae exit dauer and develop into reproductive post-dauer adults, which are in most respects indistinguishable from well-fed animals (Cassada and Russell, 1975).

In the lab, animals are maintained in an invariant environment on agar plates with abundant food. The standard laboratory wild-type strain of *C. elegans* (N2) has acquired mutations related to selection pressures of the laboratory environment (Sterken et al., 2015). These mutations have altered many behavioral phenotypes in the N2 strain including foraging, aerotaxis, and pathogen avoidance (Bendesky et al., 2011; Gray et al., 2004; Reddy et al., 2009; Rogers et al., 2006). The experience of earlylife stress inducing dauer arrest leads to changes in gene expression, endo-siRNA pathways, and altered responses to dauer pheromone in post-dauer N2 animals (Hall et al., 2013, 2010; Sims et al., 2016).

Foraging for food is an essential behavior in all animals and is directly regulated by the resource levels in the environment. Charnov’s Marginal Value Theorem models foraging behavior in terms of patch leaving events, where animals integrate internal and external cues to determine the optimal time spent exploiting the current food patch (Charnov, 1976). Exploration outside a food patch is energetically expensive and may be associated with potential risks. Thus, the animal’s behavior represents its assessment of its current patch, as well as its beliefs about the distribution of risks and resources in its broader environment. These beliefs are encoded in the functional properties of the nervous system and include both genetic predispositions and information acquired through experience.

Charnov’s framework has been used previously to study patch-leaving behavior in different *C. elegans* strains, where it was found that polymorphisms in two genes, *npr-1* and *tyra-3,* result in low leaving frequency in the N2 strain compared to the wild isolate CB4856 (HW) (Bendesky et al., 2011). These genetic differences may reflect the selective pressures of the environments inhabited by these strains, and thus represent long-term encoding of foraging strategies. In contrast with genetically determined foraging strategies, *C. elegans* foraging decisions are also plastic on the time scales of minutes and hours. For example, upon removal of food, animals first search a local area, then transition to a wider-ranging global search mode if none is found, a shortterm form of plasticity that takes place over minutes (Calhoun et al., 2014; Gray et al., 2005). *C. elegans* also acquires information about the the spatial distribution of food on the time course of hours and will adjust search strategies accordingly (Calhoun et al., 2016).

Between these time scales—the micro-evolutionary difference between strains and the short-term changes within individuals—we wondered if early-life experience might produce life-long changes in foraging strategies between genetically identical individuals. We therefore explored if early starvation stress affects foraging decisions in adults in the both the lab adapted N2 strain and the HW wild isolate. We found that early life starvation leads to permanent changes in adult foraging behavior in HW and other exploratory wild isolates, but not in the domesticated N2 strain. We identified changes in patch exploration decisions as well as long term temporal patterns in food search behavior. The *C. elegans* foraging circuitry is a large network of neurons including sensory, interneuron and pre-motor command neurons affecting navigation. We identified changes in key components of this navigation circuitry in post-dauer HW animals. Our results identify long term adaptations to early life stress in an ecologically relevant behavior and its underlying neural circuitry in *C. elegans* and suggest that such changes are favored in genetic backgrounds tuned to flexible environments.

## Results

### Early life starvation tunes adult foraging behavior in wild C. elegans

Foraging is a well-studied behavior in *C. elegans.* Under laboratory conditions, animals placed on a patch of *E. coli* periodically leave the bacterial lawn to explore their surroundings. Lawn-leaving frequency can be modulated by many external factors. For example, depleting lawns, less-preferred food, or high CO_2_ or O_2_ concentrations lead to increased leaving (Milward et al., 2011; Shtonda and Avery, 2006). The genetic background of the strain also affects foraging, with different wild-type strains having different rates of lawn leaving. Of those tested, the lab-adapted strain N2 has the lowest leaving frequency, whereas the wild isolate CB4856 (HW) is the most exploratory (Bendesky et al., 2011; Gloria-Soria and Azevedo, 2008).

Foraging can be viewed as a decision-making process that balances the animal’s current state and surroundings against beliefs about the distribution of resources and risks in the environment. We reasoned that passage through starvation-induced dauer might be an instructive cue indicating a potentially resource-poor environment, and that this should inform the beliefs used to make foraging decisions. We hypothesized that the experience of early life starvation stress and subsequent passage through the dauer stage should alter adult foraging behavior in ways consistent with an environment in which food is rare.

We placed animals on a standardized lawn and monitored their lawn-leaving behavior for one hour. We used both N2 and HW strains for this assay and compared young adult post-dauer animals, which had passed through starvation-induced dauer as larvae but had been fed ad libitum to adulthood for 48 hours prior to the assay, with “well-fed” young adult controls, which had been fed ad libitum since hatching (Figure 1A). Consistent with previous observations, the HW well-fed animals had more leaving events compared to N2 well-fed animals (Figure 1 B) (Bendesky et al., 2011; Gloria-Soria and Azevedo, 2008). Compared to the characteristic high lawn-leaving shown by the well-fed animals, the HW post-dauers had fewer leaving events. There was no difference between well-fed and post-dauer in the N2 strain. Thus, early life experience of starvation and passage through the dauer stage alters adult food seeking behavior, and this is dependent on the genetic background of the strain.

**Figure 1.**
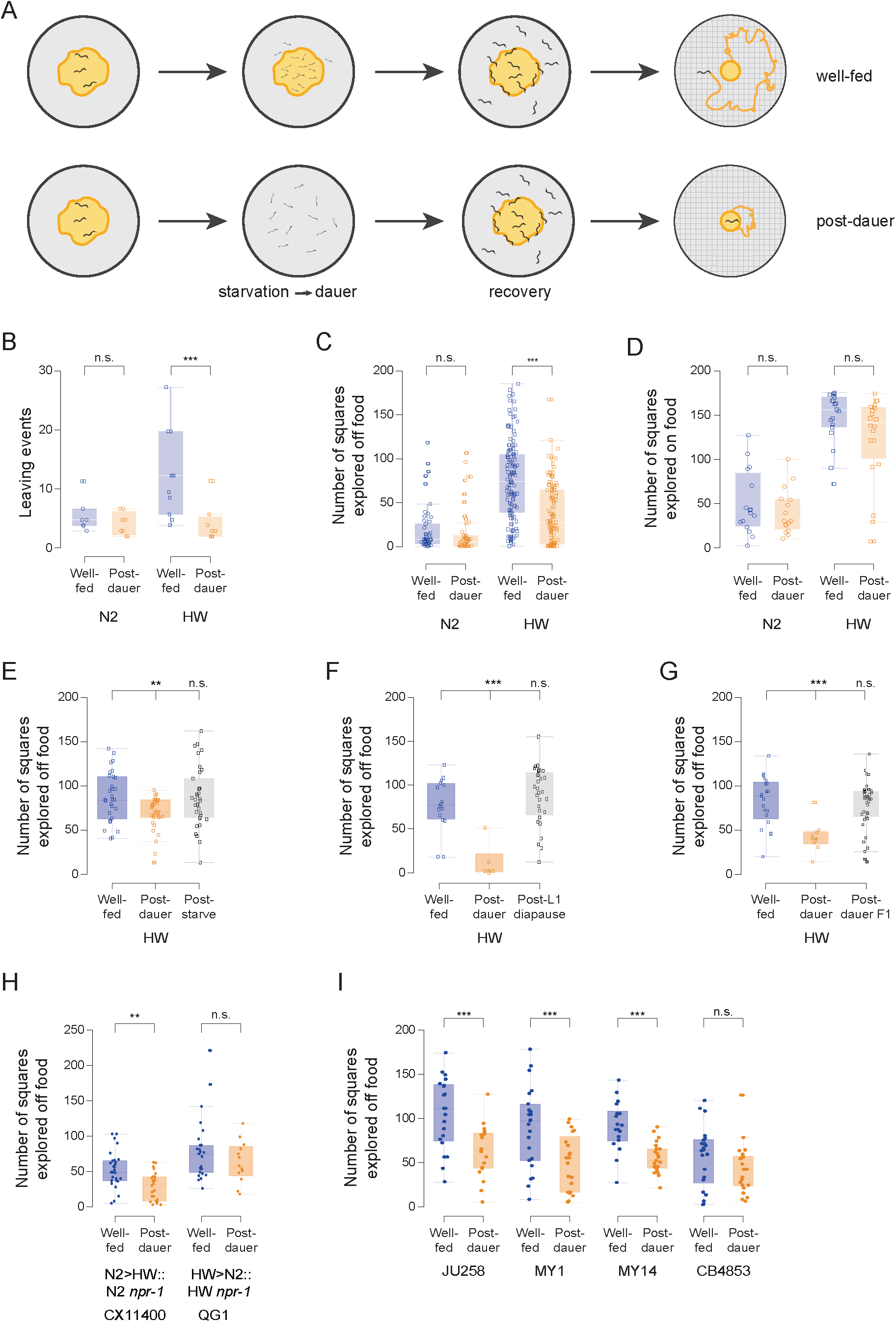
Passage through starvation-induced dauer alters adult foraging behaviour. (A) Diagrammatic representation of the experimental setup. Animals were maintained on a plate for 7 days until they starved, and a fraction of the population entered dauer. Dauers were isolated with 1% SDS, plated on seeded plates and tested 2 days later in the day 1 adult stage (post-dauer) with stage-matched well-fed controls. (B) Number of leaving events in a 1-hour assay period, each data point represents 1 assay plate containing 10 animals. Two-way ANOVA F_3,30_ = 6.9527, *P* = 0.0011, significant main effects of strain and treatment and significant interaction, \*\*\**P* < 0.001 post-hoc Student’s t-test (N2: well-fed n = 7, post-dauer n = 9; HW: well-fed n = 11, post-dauer n = 8). (C) The area explored by single animals on a plate containing a small food lawn over a 1-hour assay period. Two-way ANOVA F_3,380_= 56.5112, *P* < 0.0001, significant main effects of strain and treatment and significant interaction, \*\*\**P* < 0.001 post-hoc Student’s t-test (N2: well-fed n = 50, post-dauer n = 50; HW: well-fed n = 151, post-dauer n = 133). (D) The area explored by single animals on a plate completely covered by a food lawn over a 1-hour assay period. Two-way ANOVA F_3,67_= 37.1160, *P* < 0.0001, significant effect of strain, no effect of treatment, no interaction (N2: well-fed n = 16, post-dauer n = 16; HW: well-fed n = 19, post-dauer n = 20). (E) The same assay, comparing well-fed and post-dauer animals with animals that experienced starvation but did not enter dauer. ANOVA F_2,94_= 4.6508, *P* = 0.0119, \*\**P* < 0.01 posthoc Student’s t-test (well-fed n = 30, post-dauer n = 32, post-starvation n = 35). (G) The same assay, comparing well-fed and post-dauer animals with animals who passed through L1 diapause but not dauer. ANOVA F_2,48_= 15.1511, *P* < 0.0001, \*\**P* < 0.001 post-hoc Student’s t-test (well-fed n = 16, post-dauer n = 6, post-starvation n = 29). (G) The same assay, comparing well-fed and post-dauer animals with the F1 progeny of post-dauer animals. ANOVA F_2,68_= 9.1416, *P* = 0.0003, \*\**P* < 0.001 post-hoc Student’s t-test (well-fed n = 21, post-dauer n = 10, post-starvation n = 40). (H) The same assay comparing introgression lines where the *npr-1* locus has been swapped between N2 and HW backgrounds. Two-way ANOVA F_3,89_ = 11.2228, *P* < 0.0001, significant main effects of strain and treatment, no significant interaction, \*\**P* < 0.01 post-hoc Student’s t-test (N2>HW: well-fed n = 29, post-dauer n = 24; HW: well-fed n = 27, post-dauer n = 13). (I) The same assay testing wild strains for differences between well-fed and post-dauer. \*\*\**P* < 0.001 Student’s t-test (JU258 n = 21,19; MY1 n = 23,21; MY14 n = 18,23; CB4853 n = 22,22).

Even within isogenic *C. elegans* populations, individual variation in behavior has been reported, especially in the context of locomotory states and exploration (Stern et al., 2017). In our initial lawn-leaving assays, a large fraction of the animals never exited the lawn in the one-hour period, and the differences we observed were driven by a subpopulation of highly explorative individuals. We thus conducted single worm lawn-leaving assays in which we used a thinner lawn of bacteria to encourage more lawn leaving behavior (Milward et al., 2011). We placed single animals on standardized small lawns and left them undisturbed for one hour, after which the animals were removed, and the plates incubated in a 37°C humidified incubator overnight. *C. elegans* shed bacteria from their cuticle and defecate once every 50 seconds (Croll and Smith, 1978; Liu and Thomas, 1994), which results in a trail of bacterial colonies outside the lawn. The sinusoidal tracks left on the agar surface combined with the defecation trail allows for an easy method of tracing out each worm’s tracks as they forage outside the lawn. A grid (adapted from (Flavell et al., 2013)) was used to quantify the area explored by a single animal outside the lawn. Confirming our previous results, we found that there was no difference between N2 well-fed and post-dauer animals. However, the HW post-dauers explored significantly less area than their well-fed counterparts (Figure 1C).

We next asked if the reduced exploration behavior shown by HW post-dauers was due to gross differences in locomotion in post-dauer animals rather than foraging behavior per se. We placed animals on plates completely covered with a lawn of bacteria and measured the area explored by each animal. When on food, *C. elegans* switch between two exploration states: dwelling, characterized by low speed and a higher number of turns, and roaming, characterized by high speed and infrequent turns (Ben Arous et al., 2009; Flavell et al., 2013; Fujiwara et al., 2002). We did not find a significant difference between post-dauer and well-fed exploration in either strain while on food, indicating that post-dauer animals are not defective in locomotion but instead make different foraging decisions with respect to lawn-leaving and the extent of off-food exploration (Figure 1D).

### Dauer is necessary for long term plasticity in foraging behavior

When exposed to stressful situations like starvation, increased pheromone concentrations, or high temperature, dauer is induced in some but not all late L1 animals. We asked if transient early life starvation alone was sufficient to change adult foraging behavior. To address this question, we allowed animals to starve on plates until some of the population entered dauer. We separated the dauers from L2 animals which had not entered dauer (referred to as “post-starvation”), recovered both on food, and raised them to adulthood. These two groups experienced starvation for the same amount of time in the same developmental window. The post-starvation animals of the HW strain did not show the reduced exploration of post-dauer animals (Figure 1E).

When *C. elegans* eggs hatch in the absence of food, they go through different developmental arrest called the L1 diapause. Animals experiencing L1 diapause have been shown to have increased lifespan, which can be transgenerationally inherited for up to 3 generations (Rechavi et al., 2014) and have reduced body size, lower brood size, and altered stress responses (Jobson et al., 2015). To ask if animals experiencing the L1 diapause also show changes in foraging behavior as adults, we recovered L1 arrested animals on food till adulthood and found their foraging to be similar to that of well-fed HW animals (Figure 1F). These results indicate that passage through dauer is necessary for the long-term plasticity in foraging behavior that we observed in HW post-dauer animals. Finally, we tested whether post-dauer changes in foraging could be transgenerationally transmitted. We examined the behavior of the F1 progeny of post-dauer animals. These animals did not inherit the reduced exploration shown by their post-dauer parents (Figure 1G).

### Long-term plasticity in foraging behavior does not depend on NPR-1

Our results indicate that post-dauer animals have reduced foraging behavior in the HW wild isolate, but not in the lab adapted strain, N2. The N2 strain was maintained as a continuously reproducing population for almost two decades before it was frozen as a stock, and it has mutations acquired either through drift or adaptation to the lab environment (Sterken et al., 2015). A polymorphism in the neuropeptide receptor gene *npr-1* is the most well studied among these. The N2 strain has a gain-of-function NPR-1 receptor that results in modified aerotaxis and solitary feeding (de Bono and Bargmann, 1998). The *npr-1* polymorphism affects a range of related behaviors, including speed on food, oxygen sensing, and lawn-leaving rates (Bend-esky et al., 2011; Persson et al., 2009).

To test if the absence of behavioral plasticity in the N2 strain is an effect of the high activity *npr-1* allele, we conducted our foraging assays on two recombinant inbred advanced intercross lines (RIAILs) which had either the N2 *npr-1* locus in a HW background or vice versa (Bendesky et al., 2011). We found that the strain with N2 NPR-1 in a HW background retained the post-dauer plasticity, whereas there was no difference between well-fed and post-dauer animals in the strain with the HW NPR-1 in a N2 background (Figure 1H). This result suggests that the lack of plasticity in the N2 strain is not dependent on NPR-1. We also tested four other wild isolates of *C. elegans* known to have different lawn-leaving propensities (Bendesky et al., 2011) and found that three of them had reduced foraging behavior in post-dauer animals (Figure 1I). Thus, dauer-dependent plasticity in adult foraging behavior is common in wild strains.

### Post-dauers have reduced dispersal behavior

Exploratory behavior requires first leaving the food lawn followed by dispersing away from the lawn border. Previous work found that exploratory strains with the lower-function *npr-1* allele like the CB4856 HW strain have higher dispersal propensity than solitary strains like N2 (Gloria-Soria and Azevedo, 2008). By superimposing concentric circles of increasing diameters around the food patch, we measured how far from the original food patch the animals explored, in combination with our lawn-leaving assay (Figure 2A). We found that post-dauer animals when exploring outside the bacterial lawn did not disperse as far from the food patch, and the differences between the well-fed and post-dauer foraging were largest at circles farthest away from the lawn (Figure 2B). This indicates that post-dauer HW animals again behave like the solitary N2 strain and show reduced dispersal, in contrast to the high dispersal HW strain.

**Figure 2.**
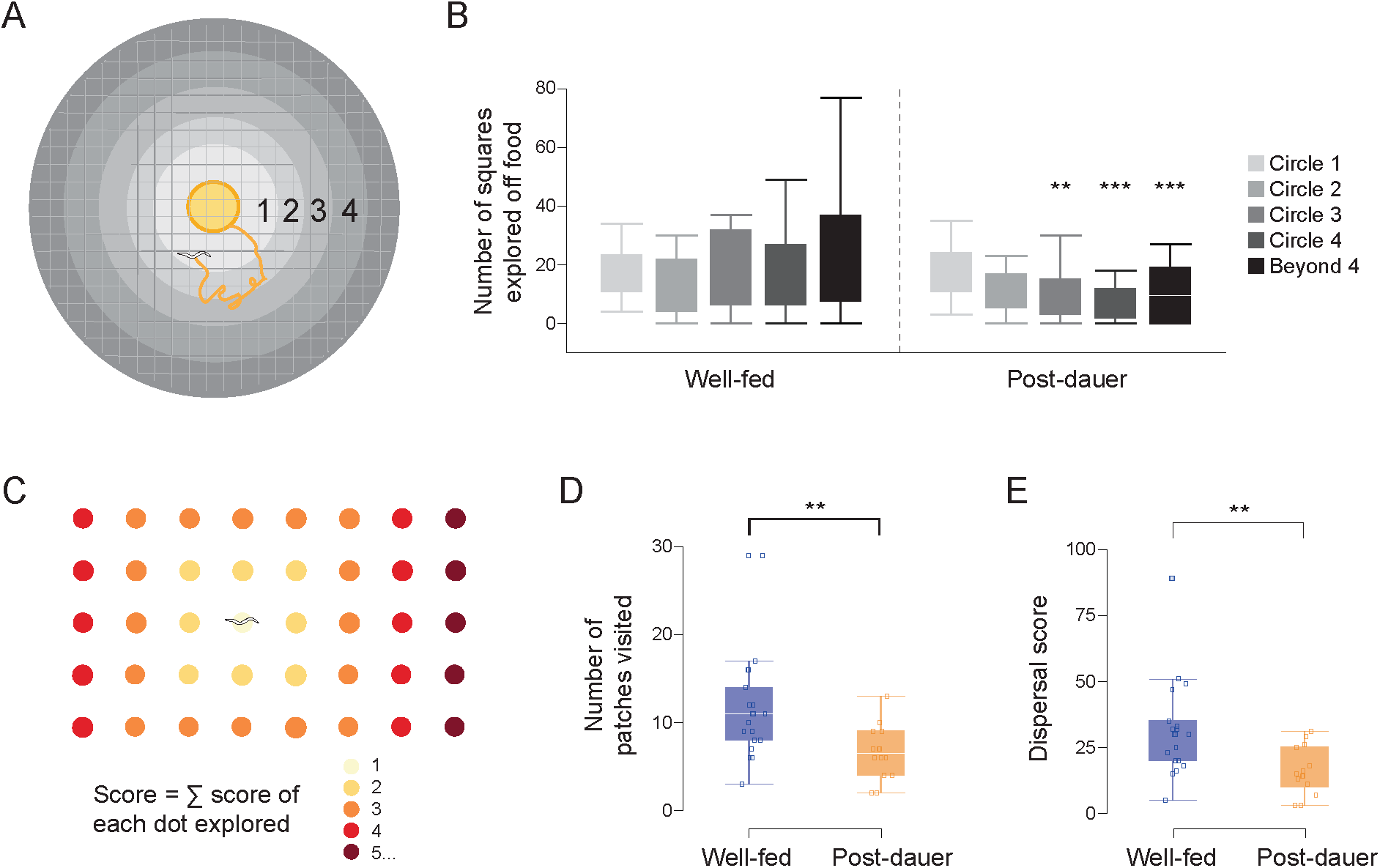
Post-dauer animals show limited dispersal range independent of food distribution. (A) Plate divided into concentric rings to estimate dispersal away from a single food source. (B) Area (squares) explored at different distances from the lawn compared by repeated measures ANOVA F_4,37_ = 5.2434, *P* = 0.0019, significant interaction between genotype and distance. \*\*\**P* < 0.001, \*\**P* < 0.01 post-hoc Student’s t-test (well-fed n = 24, post-dauer n = 18). (C) Animals explored an environment with small, evenly-distributed food patches and the number of patches visited by each individual was recorded. Animals received a weighted dispersal scoret based on the distance of each patch from the starting patch. (D) Quantification of the number of patches explored and (E) the dispersal score. \*\**P* = 0.001 Student’s t-test (well-fed n = 19, post-dauer n = 14).

In the wild, resource availability is not continuous, and animals need to travel between food patches. In contrast, our assays thus far consist of a single-patch environment. We wondered if rewarding exploratory behavior by distributing food throughout the environment might induce more exploration or modulate foraging dynamics. To test this, we arrayed a grid of tiny bacterial lawns to mimic a patchy environment (Iwanir et al., 2016) (Figure 2C). We quantified number of patches explored by a single animal over a one-hour period and found that well-fed animals explored more patches (Figure 2D). We also scored the spatial extent of exploration by using a weighted score in which the starting patch was assigned a score of 1, the nearest neighbor patch a score of 2, and so forth, with patches further away from the origin having higher scores. The dispersal score was measured as a summation of scores of all patches explored by an animal. We found that post dauer HWs explored fewer patches compared to well-fed animals and had a lower dispersal score (Figure 2E). Thus, in addition to reduced lawn-leaving and less area explored, HW post-dauers conduct more restricted searches even when food is frequently encountered.

### Post-dauers are locked in a local search mode

*C. elegans* move on solid surfaces like agar plates using an undulatory sinusoidal motion with body bends in the dorsoventral plane. Locomotion can broadly be divided into forward locomotion, reversals, and turns (Gray et al., 2005). Animals use a combination of these to conduct a biased random walk similar to bacterial chemotaxis to navigate their environment (Pierce-Shimomura et al., 1999). When placed in an environment without food, *C. elegans* have robust temporal dynamics in their food-search strategies (Calhoun et al., 2016, 2014; Gray et al., 2005; Hills et al., 2004). Immediately after removal from food, animals make short forward runs followed by frequent reversals and turns for the first 10-12 minutes. These frequent reorientations limit the total area explored, so this phase has been characterized as a “local search.” A transition to a “global search” phase follows, in which reorientations are suppressed, forward crawling bouts extended, and the animal explores a larger area.

We hypothesized that the smaller area explored off the food lawn by post-dauers may in part be an effect of high frequency of reorientations characteristic of the local search phase. We removed animals from food and video recorded them exploring an agar plate containing no food for 20 minutes (Figure 3A). To quantify foraging behavior, we calculated a path angle, a measure of the shape of the animal’s trajectory, at each time point (Figures 3B). Low path angles represent straight forward paths over short time scales, intermediate values correspond to curved paths, and high values indicate sharp turns and reversals (Figure 3C, 3D). Overall, off-food locomotion is dominated by straight forward runs (Figure 3F). We observed the previously reported (Gray et al., 2005) transition of well-fed animals from local search to global search, where reversals become less frequent after approximately 10 minutes (Figure 3E). In contrast, post-dauer animals maintained reversal frequencies characteristic of the local search phase and did not transition to global search (Figures 3D, G). The overall path angle distribution was similar for well-fed and post-dauer animals during the local search phase but showed marked differences across the distribution during global search (Figure 3F). Along with changes in reversal probability, post-dauers spend less time performing straight (low path angle) runs and more time following curved trajectories compared to well-fed animals (Figure 3H). Together, these results show that post-dauer animals do not transition to global search and are locked in a foraging mode characterized by frequent reversals and curved forward trajectories, exploring less area as a result.

**Figure 3.**
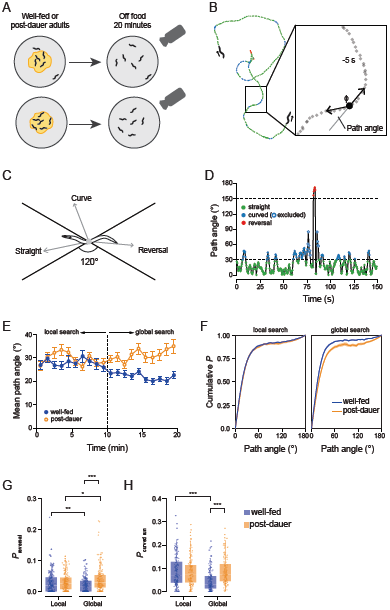
Post-dauer animals perform persistent local search. (A) Well-fed or post-dauer young adult animals were transferred in groups of 5 to plates containing no food and tracked for 20 minutes. (B) A sample path taken by one animal off food spanning 150 seconds. Foraging dynamics were analyzed using a single measure of path geometry, the angle formed by the midpoint and endpoints of a sliding 10 second time window. For clarity, path angle is defined as 180 - Φ, such that a straight path is ∼0° and a reversal is ∼180°. (C) Locomotion state definitions based on path angle. “Straight” crawling consists of forward movement within a 60° arc (path angles % 30°). Reversals are path angles > 150°. Intermediate path angles correspond to forward curved locomotion. (D) Path angle measurements over time for the example path shown in B, with color coding for locomotion state. Spurious “curved” measurements that occur from adjacent high (reversal) and low (straight) path angles are excluded. (E) Reversal probability was calculated for each track over 1-minute bins, and sample means are shown with standard error bars. (F) A comparison of mean cumulative path angle distributions for each track between well-fed and post-dauer animals during local and global search. Well-fed and post-dauer animals are similar during the local search phase. Post-dauer animals exhibit more reversals and more path curvature during the global search phase, while well-fed animals take straighter paths. (G) Sharp turn probability is dependent on prior experience and on search phase (ANOVA F_3,678_ = 20.2864, *P* < 0.0001). Well-fed and post-dauer animals were similar local search, but differed during global search, where well-fed animals reduced sharp turns. (H) Curved runs increased similarly to sharp turns, with a reduction in well-fed animals during global search (ANOVA F_3,678_ = 22.2876, *P* < 0.0001). Sample sizes vary over the time course of the experiment due to occasional tracking failures, collisions, or animals approaching the edge of the arena. The number of animals tracked at each time point ranged from n = 51–75 for well-fed and n = 53–74 for post-dauer. Post-hoc Student’s t-test \**P* < 0.05, \*\**P* < 0.01, \*\*\**P* < 0.001.

### Reversals at the lawn border and curved trajectories off-food limit post-dauer exploration

Animals never encounter food during the local-to-global search transition. However, in our initial observations, animals can exit and reenter the lawn. To understand the combined dynamics of lawn leaving and off-food exploration, we video recorded individual animals on plates consisting of a thin bacterial law (∼9 mm diameter) and observed their behavior for one hour (Figure 4A). Well-fed and post-dauer animals both tended to occupy the outer regions of the lawn, a well-defined HW behavior called bordering (Figure 4B, 4C) (de Bono and Bargmann, 1998). As in our initial assays, well-fed animals spent more time exploring off-food (Figure 4D). We noted that the mean speed and path angle of animals varied in different regions of the plate, and we used this to define four distinct behavioral zones (Figure 4E). Path angles peaked at the very edge of the lawn but were high near the border and immediately off the lawn. This likely reflects dwelling behavior while feeding, avoidance behaviors when encountering the lawn border, and local search immediately after leaving the lawn. Post-dauer animals showed higher reversals than well-fed at the lawn border, consistent with their lower rates of lawn leaving (Figure 4F). We were surprised to see that post-dauer animals did not show any enhanced reversals off lawn compared to well-fed. Instead we observed a striking difference in the amount of time spent following curved paths (Figure 4G). These are forward runs that deviate systematically from a straight path. Intuitively, curved paths may both limit search area and tend to return an animal to the lawn, limiting off-food exploration.

**Figure 4.**
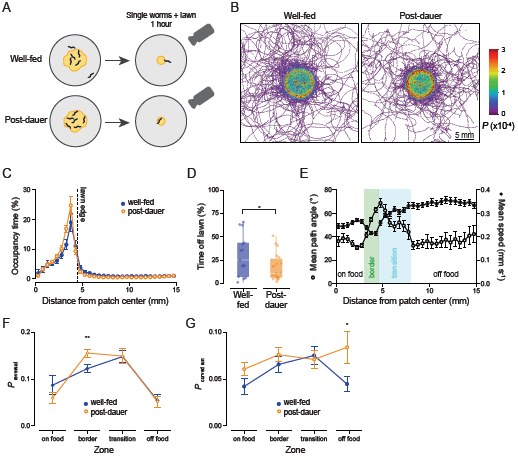
Locomotion differences underlying lawn-leaving and exploration near food patches. (A) Well-fed or post-dauer young adult animals were transferred individually to agar plates containing a small lawn and video recorded for 1 hour. (B) Heat map of occupancy probability of each point. (C) Mean percent of time spent by distance from the center of the food patch in 0.5 mm bins. (D) Average percent of time spent off lawn by well-fed and post-dauer animals. (E) Locomotion variables (path angle and speed) as a function of distance from path center across all animals were used to define zones for subsequent analyses. (F) Sharp turn probability as a function of zone for well-fed and post-dauer animals (H) Curved run probability as a function of zone for well-fed and post-dauer animals. Well-fed n = 15, post-dauer n = 20. \**P* < 0.05, \*\**P* < 0.01, Student’s t-test.

### Post-dauer animals show altered activity in a navigation circuit

We next aimed to identify the neural correlates of this long-term behavioral change in post-dauer HW animals. Foraging and lawn-leaving are multi-modal behaviors with dozens of neurons implicated in their regulation. Our behavioral assays indicated that post-dauer animals have higher number of reorientations and turns at both the lawn boundary as well as off food environments after the local search phase (Figure 3G, Figure 4F). We thus decided to focus on neurons involved in initiating reversals and hypothesized that they may be differentially regulated by food cues in well-fed and post-dauer animals. Even in response to strong stimuli, reversal responses are probabilistic. Previous work showed that a circuit of two interneurons, AIB and RIM, and the pre-motor command neuron AVA regulate stimulus-evoked reversal probability (Gordus et al., 2015). Activity in AVA is directly linked to spontaneous reversal initiation in freely behaving animals (Chalfie et al., 1985; Gray et al., 2005; Guo et al., 2009). AIB acts as a first order interneuron receiving input from multiple food-sensing neurons and contributes to deterministic reversal output in response to removal of attractive odors; in contrast, RIM activation is associated with increased variability in behavioral output (Gordus et al., 2015). AIB and RIM are also important in learning paradigms where tyraminergic inputs from RIM regulate long term memory of pathogen avoidance (Jin et al., 2016).

Calcium imaging in *C. elegans* is often performed in microfluidic devices by presenting a large, discrete change in sensory input such as step changes between a odorant and a stimulus-free buffer solution. In contrast, the area around a bacterial lawn border is a transitional region where mechanical properties, levels of CO_2_ and O_2_, and various food-related tastants and odorants are diffusing and distributed in overlapping and complex ways. While we cannot recapitulate all features of a lawn border, we chose to use higher and lower concentrations of a bacterial food culture as competing stimuli rather than an “all or none” stimulus. Our aim was to produce a multi-modal sensory experience that is less extreme than sudden removal of all food-related cues and that may better approximate the differences experienced around a lawn border.

To avoid short-term starvation effects, animals were removed from plates and placed immediately in the higher concentration food culture and maintained in this medium while being immobilized in a microfluidic device (Chronis et al., 2007). They were then imaged for 1 minute, followed by 1-minute exposure to a lower food concentration, then 1 minute again in the higher concentration.

AVA, AIB, and RIM have bimodal high (ON) and low activity (OFF) states, and transitions between states can be odor-driven or spontaneous (Gordus et al., 2015). Following these authors, we classified ON and OFF states for each neuron (Figure 5A). Not surprisingly, all three neurons showed much more variability and ambiguous responses in response to changes in food levels compared to the more deterministic responses previously seen in response to the presence or absence of a pure odor (Figure 5B-D) (Gordus et al., 2015). While we did not see consistent differences in states averaged across all animals, we observed that AVA and RIM have higher numbers of state transitions in postauer animals (Figure 5B, D). We did not find any difference in AIB. This suggests that in same stimulus context, post-dauer animals generate more variable responses and more dynamic state changes over short time scales compared to well-fed animals.

**Figure 5.**
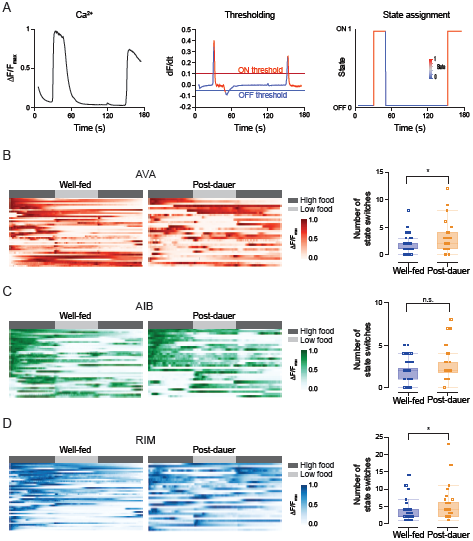
State dynamics are altered in navigation circuitry in post-dauer animals. (A) Classification of ON/OFF states for AVA, AIB and RIM. Representative normalized trace (left) showing rise and fall of GCaMP intensity; time derivative (dF/dt) of the same trace (center) with thresholds for ON and OFF states for each neuron; ON and OFF state representation of the same trace (right). See Methods for details. (B) Heat maps of AVA activity in well-fed and post-dauer animals (left). Comparison of the number of state switches (right). (C) Heat maps of AIB activity in well-fed and post-dauer animals (left). Comparison of the number of state switches (right). (D) Heat maps of RIM activity in well-fed and post-dauer animals (left). Comparison of the number of state switches (right). All heat maps are ordered by the number of state switches. Each row represents a single animal, dark grey bars indicate high food stimulus, light grey indicates low food dilutions. \**P* < 0.05 Wilcoxon test, well-fed n = 42, post-dauer n = 30.

### Silencing candidate interneurons reduces exploration

Because we observed more dynamic activity in AVA and RIM in post-dauer animals, we hypothesized that silencing these neurons might increase foraging to a more well-fed-like state, likely through preventing reversals. To do so, we used the histamine-gated chloride channel of *Drosophila melanogaster* (HisCl1) under neuron-specific promoters, which allows neurons to be selectively inhibited in the presence of exogenous histamine (Pokala et al., 2014).

We tested the off-food foraging behavior of animals expressing HisCl1 individually in AVA, AIB or RIM in the presence and absence of histamine using the same assay used previously (Figure 1A, 1C). We expected that suppressing reversals by inhibiting this circuit would cause animals to show longer forward runs and increase the area explored during foraging. Surprisingly, silencing either AVA or RIM with HisCl1 dramatically decreased foraging area in well-fed animals and did not increase exploration in post-dauers. Silencing AIB had no effect on foraging area (Figure 6A). Previous work has reported HisCl1 silencing of AVA can also decrease speed of forward locomotion in addition to decreasing reversals (Pokala et al., 2014). The lower foraging in well-fed animals in our lawn-leaving assay may be partially explained by this reduction in speed.

**Figure 6.**
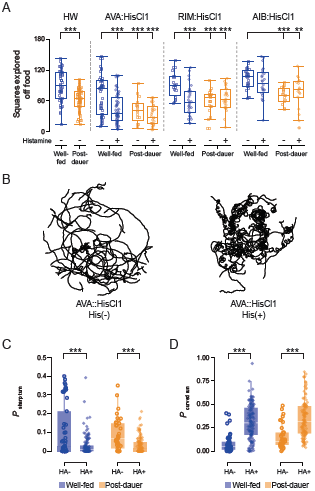
Suppression of the reversal circuit limits foraging area by causing curved crawling. (A) Area explored by animals expressing the histamine-gated chloride channel HisCl1 in AVA, AIB, or RIM in the absence or presence of histamine. Two-way ANOVAs were significant for all, with significant main effects of histamine treatment and well-fed/post-dauer, for AVA and RIM. AIB was significant for well-fed/post-dauer but not for histamine treatment. The interaction was only significant for RIM. AVA: F_3,153_ = 13.0456, *P* < 0.0001; AIB: F_3,81_ = 6.8588, *P* = 0.0004; RIM: F_3,92_ = 7.5422, *P* = 0.0001.\*\*\**P* < 0.001 post-hoc Student’s t-test comparing theeffect of histamine for each group. n (left to right) = 50, 68, 49, 59, 17, 32, 26, 24, 16, 19, 20, 29, 21, 26.Example traces of trajectories of AVA::HisCl1 animals in the presence and absence of histamine.Quantitation (as in Figure 3) of reversals for animals expressing HisCl1 in AVA in the presence and absence of histamine. ANOVA F_3,349_= 20.6722, *P* < 0.0001, \*\*\**P* < 0.001 post-hoc Student’s t-test.Quantitation (as in Figure 3) of reversals for animals expressing HisCl1 in AVA in the presence and absence of histamine. ANOVA F_3,349_= 34.8069, *P* < 0.0001, \*\*\**P* < 0.001 post-hoc Student’s t-test.

In addition to increased sharp turns at lawn boundaries (Figure 4F), our behavioral experiments also showed that lower area explored during foraging by post-dauers is in part caused by increased number of curved runs in an off-food environment (Figure 3H, 4G). To examine the causes of decreased search in detail, we performed off-food tracking experiments as in Figure 3. We observed the AVA::HisCl1 animals showed highly curved trajectories in the presence of histamine (Figure 6B). Suppression of AVA did indeed prevent reversals (Figure 6C), but this was accompanied with a large increase in the proportion of time spent in curved forward runs (Figure 6D). It is this increase in path curvature that causes a reduction in area explored. This result suggests that manipulating the reversal command neuron AVA can have dramatic effects on other aspects of locomotion. We also observed increased path curvature, albeit less dramatically, in post-dauer animals exploring off-food in our lawn-leaving assays (Figure 4). Thus, persistently curved locomotion is a behavioral strategy that limits search area, both in addition to or independently from reversal frequency, and these two behaviors may share common circuitry.

## Discussion

The environment can have profound effects on development, leading to permanent changes in physiology and behaviour. In combination with genetic predispositions, such environmental factors can be major determinants of adult phenotypes. Nutritional and other stressors during pre- and perinatal development are important risk factors for adult disease in humans. Researchers have theorized that some diseases, ranging from type 2 diabetes to behavioural and affective disorders, arise from adaptive forms of developmental plasticity that have evolved to optimize or tune traits to a predicted adult environment (Byrne and Phillips, 2000; Sih, 2011). Mismatches between the predicted and actual environment lead to detrimental health conditions. There is a growing appreciation for the role of developmental stress in adult behavioural traits and in mental disorders, including schizophrenia, depression, anxiety, and sleep disorders (Bale et al., 2010; Datta et al., 2000; Hulshoff Pol et al., 2000; Piper et al., 2012; Verdoux, 2004). While many observations support the developmental origins theory (Barker, 2004), the relevant mechanisms remain obscure due to the complexity of gene-environment interactions and the long delays between developmental cause and adult effect.

In *C. elegans,* dauer is a well-characterized response to environmental stress, and post-dauer animals show long-term changes in gene expression and sensory neuroanatomy and physiology (Golden and Riddle, 1984; Hall et al., 2010; Sims et al., 2016). Here, we show that starvation stress also acts to program adult foraging behavior such that strains genetically predisposed to be highly exploratory become “cautious” foragers and resemble non-exploratory strains. We interpret this to mean that starvation stress during early-life is an indicator of a potentially resource-poor environment, and that this information biases animals toward favoring feeding even on a poor-quality food source over exploring in search of better resources.

Trait plasticity is expected to to arise in fluctuating environments. In stable environments, we expect that foraging strategies may become genetically fixed, as plasticity comes with its own costs (Houston and McNamara, 1992). The lack of foraging plasticity in N2 might represent adaptation to the lab environment, in which animals are raised on single-lawn plates such that search off-food is never advantageous. Alternatively, it may be that less-exploratory genotypes can exhibit plasticity, but that it is difficult to measure further reductions from already low foraging rates.

We observed two major differences in foraging between well-fed and post-dauer animals. The first was that in the presence of food, post-dauer animals are much less likely to leave the lawn and explore less area off the lawn when they do so (Figure 1). This limitation in the area explored is not affected by the likelihood of finding additional food resources (Figure 2). Limiting search area in *C. elegans* is usually accomplished by increasing the frequency of reversals and limiting the duration of forward crawling (Calhoun et al., 2014; Gray et al., 2005). In contrast, we observed that the limited off-food search conducted in the presence of a food patch was not accompanied by increasing reversal probability, but by increasing the curvature of forward locomotion (Figure 3). We also observed that in the absence of food, well-fed and post-dauer animals show similar reversal probabilities during the local search phase, but over time, as well-fed animals transition to the low reversal probability global search phase, post-dauer animals stay locked in local search mode.

Both spontaneous and sensory-evoked reversals involve the command interneuron AVA, and AVA’s activity state is under complex regulation (Gordus et al., 2015). We observed that the AIB-RIM-AVA micro-circuit undergoes more frequent state transitions in post-dauer animals compared to well-fed. This observation is at least consistent with our observation that post-dauers are locked into a local search mode characterized by frequently alternating between forward and reverse crawling. Finally, chemogenetic inhibition of this network had unexpected effects on foraging dynamics. While reversals were suppressed, as expected, foraging area was dramatically limited because of highly curved crawling trajectories. Because we identified path curvature as a feature of limiting exploration in post-dauers, this suggests a compelling link between the AVA circuit and movement patterns beyond the initiation of reversals. Tightly curved crawling has been observed in mutations in the LET-60 Ras MAP Kinase pathway, where it acts upstream of glutamate receptor localization, suggesting a link between excitability in the central motor circuit and locomotion dynamics (Hamakawa et al., 2015).

The dauer stage represents an extreme case of responsiveness to developmental stress. Passage through dauer not only confers resistance to these stressors in the short-term, but as we have shown here confers indelible, adaptive behavioral traits tuned to a low-resource environment. Life-long changes in foraging strategies may arise from changes in either the structure or functional properties of the nervous system, or both. At present, our observations of changes in neural circuit activity cannot distinguish between these possibilities. However, the evolutionarily ancient origins of developmental tuning and adaptive plasiticity speaks to the ubiquity of the challenge of balancing the risks and opportunities of foraging behavior, and suggests that *C. elegans* is an attractive system in which to study the neurodevelopmental basis of this plasticity.

## Materials and Methods

### Nematode culture

*C. elegans* strains were maintained at 20–22°C on nematode growth medium (NGM) plates seeded with *E. coli* HB101 strain (Brenner, 1974). Young-adult hermaphrodites were used for all experiments. Transgenic strains originating in N2 were backcrossed to the HW strain at least 15 times and subsequent strains were behaviorally validated (bordering, clumping, dauer-dependent exploratory behavior). All replicates of behavioral experiments included matched controls (HW well-fed and post-dauer) tested in parallel. Unless otherwise indicated, all assays were done on at least two different days with three or more replicates. Calcium imaging was done on multiple days with matched controls. Strains used:

N2 Bristol, England

CB4856 Hawaii

CB4853, Altadena, California

MY1, Lingen, Germany (Haber et al., 2004)

MY14, Mecklenbeck, Germany (Haber et al., 2004)

JU258, Madeira Island, Portugal (Marie-Anne Felix)

QG1 [qgIR1 (X: 4,754,307–4,864,273 CB4856>N2) X]

CX11400 [kyIR9 (X: ∼4745910–∼4927296,

N2>CB4856) X]

AIB/AVA/RIM::GCaMP3

3, MMH108[kyEx4965>CB4856], derived from CX14496 (Gordus et al., 2015)

AIB::HisCl1, MMH110 [kyEx4866>CB4856], derived from CX14848 (Pokala et al., 2014)

AVA::HisCl1, MMH111 [kyEx4864>CB4856], derived from CX14846 (Pokala et al., 2014

RIM::HisCl1, MMH112 [kyEx5464>CB4856], derived from CX16040 (Pokala et al., 2014)

### Generation of well-fed control and post-dauer animals

Control “well-fed” animals were maintained on abundant food throughout their development on NGM plates seeded with *E. coli* HB101. Plates left for 5–7 days led to depletion of food and starvation-induced dauer arrest. Dauer larvae were isolated by incubating animals in 1% sodium dodecyl sulphate (Fisher, Saint-Laurent, Quebec) for 30 minutes, which kills non-dauer animals. Dauer larvae were washed and placed on HB101 seeded plates until they reached young adulthood (approximately 2 days) to obtain “post-dauer” animals.

For L1 diapause experiments, gravid adults were treated with sodium hypochlorite to isolate eggs, which were then put on a plate with no food. After entering diapause, these animals were picked manually and placed back on food until young adulthood to obtain “post-L1 diapause” animals. For “post-starvation” animals, we first synchronized the population by bleaching and plated isolated eggs on a seeded plate. As this synced population exhausted food over a period of 4-5 days, a fraction of them go into the dauer stage. We selected animals which were starved for an equivalent time but did not go into dauer. Non-dauer larval worms were identified by morphology, manually picked, and put back on food until young adulthood and termed “post-starvation” animals. The same plate was then treated with SDS to isolate dauers to obtain “post-dauer” animals.

### Behavioral assays

#### Lawn leaving and exploration

For the lawn-leaving assays in Figure 1, 6cm NGM plates were seeded with 10 μl of a fresh overnight culture of *E. coli* HB101 (diluted in LB to OD600 = 0.2), 24 hours before the assay. Ten young-adult animals were allowed equilibrate on the lawn for 30 minutes followed by a 60 minutes assay period. During the assay period, behavior was monitored manually by observers blinded to strains and conditions. The number of animals exploring outside the lawn was recorded at 2-minute intervals and number of leaving and return events quantified.

For quantifying area explored off food, 9 cm NGM plates were seeded with 10 μl of fresh overnight culture of *E. coli* HB101 (diluted in water to OD_600_ = 0.1), 20–24 hours before the assay. A single young-adult animal was placed on each plate for an assay period of 1 hour. The animal was removed after the assay period and the plates incubated overnight in a 37°C humidified incubator, which allows bacteria shed and egested by worms to grow into bacterial colonies, forming a path. Worm tracks were traced out manually under the microscope the next day, and the area explored quantified by superimposing the trail on a 35 mm^2^ grid. Plates where the animal crawled off the agar onto the walls of the plate and died were rejected. Temperature and humidity fluctuations in the laboratory environment affect baseline exploration rates, so the HW well-fed and post-dauers were always tested together with any other strains/conditions tested.

To measure area explored on food, 6 cm NGM plates were seeded uniformly with 600 μl of fresh overnight *E. coli* HB101 culture. A single animal was transferred to the center of the plate and allowed to move around for 16 hours. The animal was removed after the assay and its tracks on the bacterial lawn were traced and quantified with a grid, identical to the previous assay.

### Dispersal assays

Concentric circles were superimposed on the grid used previously to measure dispersal behavior. Circles were numbered from 1 to 4, with 1 being the circle closest to the central bacterial lawn and 4 being the farthest. This was done in conjunction with exploration assays and the same data were used for both analyses. For patch exploration behavior, a grid of 32 micro patches was used. 2 μl of *E. coli* HB101 culture was seeded for each patch, diluted to the same density as the exploration assays performed before. A single animal was placed upon a central food patch and the number of patches explored quantified. To further quantify the dispersal from the starting patch, a score was assigned to individual patches. The initial patch was assigned a score of 1, with its first neighbor as 2, second neighbor as 3 and so on. The summation of the scores of the patches explored by the animal was taken as a measure of its dispersal from the origin.

### Local search assays

For local search assays, animals were washed for 5 minutes in NGM buffer and picked onto plates with no food. Animals were allowed to explore for 20 minutes and videos were tracked as described below.

### Tracking

Animals were obliquely illuminated using circular strips of 626 nm LEDs (superbrightleds.com, St. Louis, Missouri) and behavior was recorded with Logitech C210 USB Webcams at 2 frames per second. Background subtraction and image processing were done with the FIJI distribution of ImageJ, and the TrackMate plugin was used to acquire XY coordinates of worm centroids in each frame (Schindelin et al., 2012; Tinevez et al., 2017). At each time point we calculated an angle, Φ = cos^−1^(*f* • *b*) where the unit vectors extend from the current position to positions 5 seconds forward or <u>b</u>ackward, respectively. Path angle was defined as 180-Φ so that, intuitively, low angles would correspond to a straight path and high angle values to reve rsals and sharp turns.

### Calcium imaging

Restrained animals were imaged in microfluidic chips (Chronis et al., 2007) with 1 minute switches of 25% (high) and 7% (low) dilutions of freshly prepared overnight cultures of *E.coli* HB101 strain grown at 28. in NGM liquid. Animals were washed in the high food dilution for 5 mins before introduction into the chip, with 1 mM tetramisole (Sigma-Aldrich, Oakville, Ontario) present in the loading syringe and channel to avoid motion artifacts for the simultaneous imaging of AVA, AIB and RIM. Previous work has shown that using tetramisole at even higher concentrations (10 mM) do not significantly affect neuronal dynamics (Gordus et al., 2015). Animals were not starved before imaging to avoid potential confound with short term starvation effects. TIFF stacks were acquired at 5 fps with 100 ms exposure using a 40X silicone-immersion objective on an Olympus IX83 inverted microscope (Olympus, Richmond Hill, Ontario). Fluid flow was controlled with a Valvebank solenoid pinch valve array (AutoMate Scientific, Berkeley, California).

GCaMP3 fluorescence intensity was measured with FIJI (ImageJ). All further analysis was done in JMP 13.0 (SAS, Toronto, Ontario). For simultaneous imaging of AVA, AIB, and RIM, only traces where all three neurons could be captured without motion artifacts were used. Neurons that showed no dynamics over the three-minute experiment were classified as OFF. Fluorescence intensity was normalized from 0 to 1 (F-F_min_)/(F_max_-F_min_). Mean normalized fluorescence values during the first 2 secs of recording was calculated. If normalized fluorescence was >0.5, the initial state was classified as ON. Next, time derivatives were calculated for each neuron individually with a lag of 2 seconds. We then determined thresholds for classifying ON and OFF states, depending on the dynamics of the neuron. AVA and AIB has comparatively fast rise and fall dynamics (Gordus et al., 2015), and dF/dt values greater than 0.1 (for a time period τ = 2 s) an assigned an ON state (OFF threshold =−0.05). RIM has slower dynamics (Gordus et al., 2015) and thus dF/dt thresholds for state assignment lower (ON threshold = 0.04, OFF threshold =−0.02, τ = 2 s). Switches from ON?>OFF or vice versa were counted as a measure of spontaneous dynamics in each neuron individually.

### Histamine experiments

For all histamine silencing behavioral assays, animals were picked onto 10 mM histamine plates seeded with the same amount of bacteria as maintenance plates (prepared a week before and stored at 4.) for 30 mins before the start of the assay. Animals were then moved to either lawn-leaving assay setups (described earlier) or plates with no food (local search assays) for experiments.

## Acknowledgements

This work was supported by funding from McGill University, the National Science and Engineering Research Council (NSERC) (RGPIN/05117-2014), the Canadian Foundation for Innovation (CFI) (32581), and the Canada Research Chairs Program (950-231541). We thank Xinyu Liu and Xianke Dong (McGill University) for providing photoresist masters for microfluidic devices, Cornelia Bargmann (Rockefeller University) for providing HisCl1 and GCaMP strains. We thank all members of the lab for helpful discussions and comments. Some strains were provided by the *Caenorhabditis* Genetics Center (CGC), which is funded by NIH Office of Research Infrastructure Programs (P40 OD010440).

## Competing Interests

The authors declare that they have no competing interests.

